# The beginning of the end: a chromosomal assembly of the New World malaria mosquito ends with a novel telomere

**DOI:** 10.1101/2020.04.17.047084

**Authors:** Austin Compton, Jiangtao Liang, Chujia Chen, Varvara Lukyanchikova, Yumin Qi, Mark Potters, Robert Settlage, Dustin Miller, Stephane Deschamps, Chunhong Mao, Victor Llaca, Igor V. Sharakhov, Zhijian Tu

## Abstract

Chromosome level assemblies are accumulating in various taxonomic groups including mosquitoes. However, even in the few reference-quality mosquito assemblies, a significant portion of the heterochromatic regions including telomeres remain unresolved. Here we produce a *de novo* assembly of the New World malaria mosquito, *Anopheles albimanus* by integrating Oxford Nanopore sequencing, Illumina, Hi-C and optical mapping. This 172.6 Mbps female assembly, which we call AalbS3, is obtained by scaffolding polished large contigs (contig N50=13.7 Mbps) into three chromosomes. All chromosome arms end with telomeric repeats, which is the first in mosquito assemblies and represents a significant step towards the completion of a genome assembly. These telomeres consist of tandem repeats of a novel 30-32 bp telomeric <u>r</u>epeat <u>u</u>nit (TRU) and are confirmed by analysing the termini of long reads and through both chromosomal *in situ* hybridization and a Bal31 sensitivity assay. The AalbS3 assembly included previously uncharacterized centromeric and rDNA clusters and more than doubled the content of transposable elements and other repetitive sequences. This telomere-to-telomere assembly, although still containing gaps, represents a significant step towards resolving biologically important but previously hidden genomic components. The comparison of different scaffolding methods will also inform future efforts to obtain reference-quality genomes for other mosquito species.

**100-word Article Summary:** We report AalbS3, a telomere-to-telomere assembly of the *Anopheles albimanus* genome produced by integrating advancing technologies including Oxford Nanopore and Bionano optical mapping. AalbS3 features much of the difficult-to-assemble genomic ‘dark matters’ including previously missed transposons, centromeres and rDNA clusters. We describe novel telomeric repeats that are confirmed by analysis of long reads and by telomere hybridization assays. This reference-quality assembly represents a significant step towards completing the genomic puzzle pieces and informs efforts to improve the assembly of other mosquito species. Future research into the relationship between telomere and mosquito life span may have significant implications to disease control.

## INTRODUCTION

Mosquitoes belonging to the genus *Anopheles* transmit malaria parasites which cause one of the most devastating diseases known to mankind. In 2015, an international consortium published genome assemblies for 16 *Anopheles* species, which represent important resources and provided significant evolutionary insights [1]. The majority of these assemblies, limited by short read technologies, remain fragmented especially in repeat-rich heterochromatic regions. Recent integration of new scaffolding methods with Pacific Biosciences (PacBio) Single Molecule Real Time (SMRT) sequencing, which produces long reads, has enabled marked improvements to a few mosquito assemblies [2-4]. However, a substantial portion of the heterochromatic regions remain unresolved even in the best mosquito assemblies. In this study, we select the New World malaria mosquito, *An. albimanus*, to explore approaches towards more inclusive mosquito genome assemblies that enable discoveries of biologically important but previously hidden genomic components. *An. albimanus* belongs to the subgenus Nyssorhynchus and is a vector of the malaria parasite *Plasmodium vivax.* This mosquito inhabits the neotropical regions of Latin America, stretching from southern United States to Peru and the Caribbean Islands [5-10]. It has one of the smallest genomes within the genus making it a suitable species for achieving our objectives. In 2017, extensive physical mapping of this genome corrected several mis-assemblies and placed contigs onto five chromosome arms, generating a significantly improved AalbS2 assembly [11]. Here we aim to produce an improved *de novo* assembly by integrating Oxford Nanopore sequencing, Illumina, Hi-C and a recent advancement in BioNano optical mapping chemistry called Direct Labelling and Staining (DLS), which results in longer molecules and produces more contiguous maps [12, 13]. The resulting assembly, AalbS3, represents a significant improvement as it includes previously uncharacterized centromeric and rDNA clusters and added 7.4 Mbps sequences, most of which are repeats. More importantly, AalbS3 is organized into three chromosomes, with all five chromosome arms ending with telomeric repeats, which is the first in mosquito assemblies and indicates high quality. The discovery of a 30-32 bp Telomeric Repeat Unit (TRU), different from any known TRUs, provides an opportunity to investigate telomere function and evolution in mosquitoes. This discovery also opens the door to investigations into the relationship between telomere length and female mosquito life span, a trait relevant to disease transmission.

## RESULTS

### Assembly of the *Anopheles albimanus* genome

We performed ONT sequencing using gDNA isolated from sibling females from a single-pair mating (Figure 1). We also sequenced both parents using Illumina for polishing. Raw ONT reads were base called using albacore and quality-filtered sequences were assembled with Canu [14] applying a read length filter, excluding those shorter than 2 kbps. This generated a 197.39 Mbps genome consisting of 660 contigs with an N50 of 13.7 Mbps. These contigs were polished with Pilon [15] using parental Illumina short reads. To scaffold these contigs, we performed optical mapping using ultra high molecular weight genomic DNA isolated from sibling males (Figure 1). Direct Label and Stain (DLS) chemistry was used for this purpose, which produced 342 Gbps of molecules with a N50 of 373 kbps. The resulting Bionano assembly included 29 maps with an N50 of 51 Mbp and a total length of 223 Mbp. Bionano maps were then aligned to the previously mentioned 197.39 Mbps sequence assembly to generate hybrid scaffolds. The alignment produced only 8 scaffolds, with a total scaffold length of 184.23 Mbp. One of the scaffolds corresponded to a 11.8 Mbp map with no or little corresponding sequence from the female sequence assembly, likely corresponding to the Y chromosome as males were used for Bionano mapping. A 5.2 Mb scaffold corresponds to the circular genome of *Serratia marcescens*, a known bacterium in the mosquito microbiome [16]. The remaining sequences were manually curated to trim sequence overlaps, remove secondary contigs due to heterozygosity, break up mis-assemblies. The curated assembly consists of five superscaffolds including the three chromosomes X, 2, and 3 (Figure 2A), a ∼803 kb rDNA cluster and an unplaced sequence of ∼238 kb. The Hi-C contact matrix of the three chromosomes (Figure 2B) indicates good quality. Two scaffolds representing centromeric regions of chromosome 2 and 3 were added to the above five Bionano superscaffolds, resulting in the final 172.6 Mbps AalbS3 assembly (Table 1, Figure 2, and Supplemental Table S1). The two centromeric sequences contain highly repetitive tandem repeats and they were produced by Hi-C scaffolding of the polished contigs produced by Canu, as described below. The completeness of the assembly was measured using a Benchmarking Universal Single Copy Orthologs (BUSCO) test [17, 18] and scored a 99.0% with 98.2% of genes represented in this genome as complete and single copy and 0.8% as complete and duplicated (Table 1 and Supplemental Table S2).

**Table 1.**
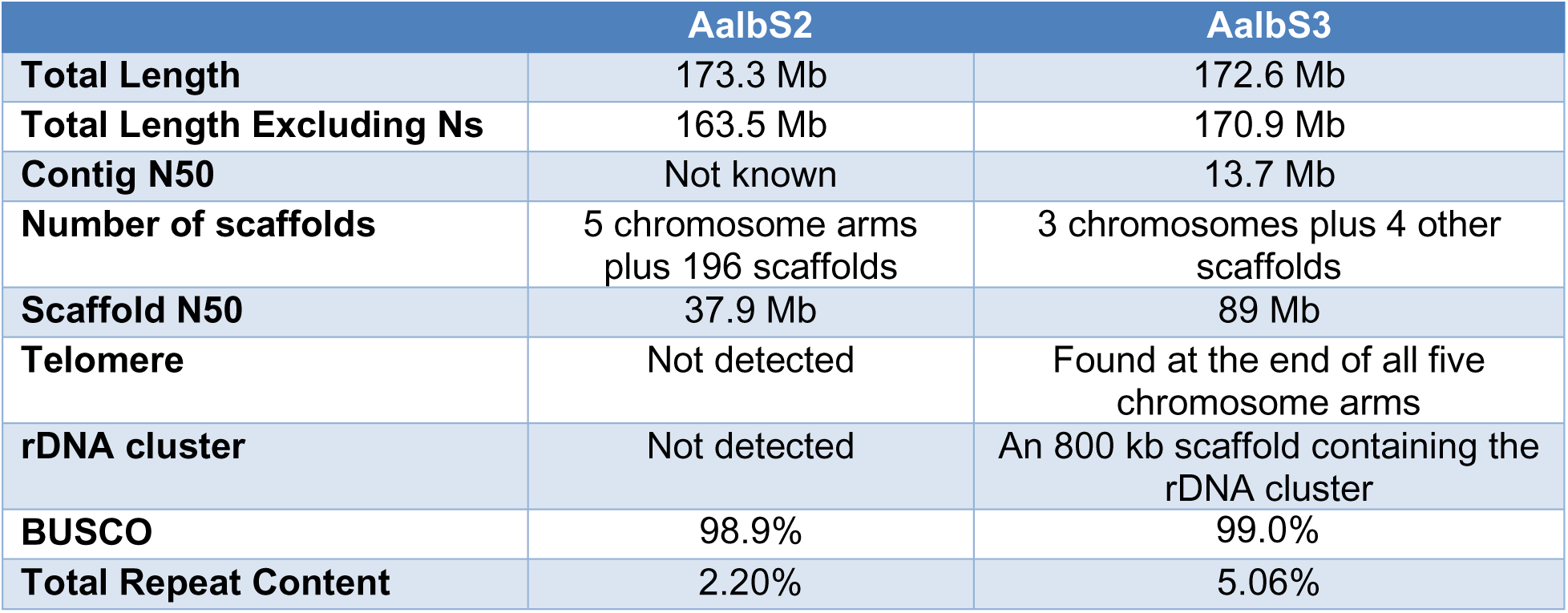
Comparison overview of AalbS2 and AalbS3.

|  | **AalbS2** | **AalbS3** |
| --- | --- | --- |
| **Total Length** | 173.3 Mb | 172.6 Mb |
| **Total Length Excluding Ns** | 163.5 Mb | 170.9 Mb |
| **Contig N50** | Not known | 13.7 Mb |
| **Number of scaffolds** | 5 chromosome arms plus 196 scaffolds | 3 chromosomes plus 4 other scaffolds |
| **Scaffold N50** | 37.9 Mb | 89 Mb |
| **Telomere** | Not detected | Found at the end of all five chromosome arms |
| **rDNA cluster** | Not detected | An 800 kb scaffold containing the rDNA cluster |
| **BUSCO** | 98.9% | 99.0% |
| **Total Repeat Content** | 2.20% | 5.06% |

**Figure 1.**
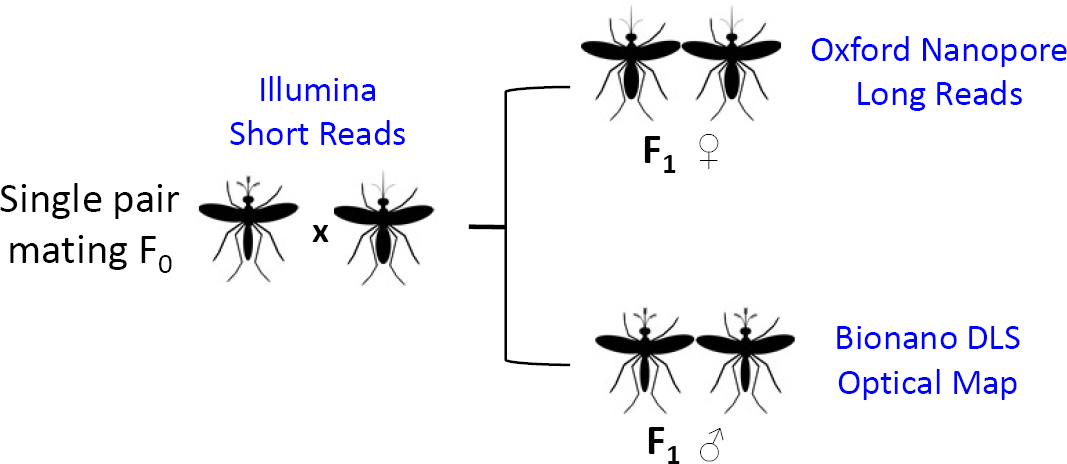
Sample and data collection scheme. The left shows the cross of a single male and female mosquito to produce male and female F1 offspring. The F0 father and mother were sequenced individually using Illumina HiSeq to obtain short reads. Genomic DNA from a pool of the female F1 sibling was sequenced using three Oxford Nanopore MinION flow cells to produce long reads. Genomic DNA from a pool of the male F1 sibling was used to generate Bionano DLS optical mapping. Oxford Nanopore reads were used to generate the contigs using Canu (ref) and the Canu contigs were polished using the parental Illumina reads. The polished Canu contigs were scaffolded using the Bionano DLS optical map to generate the AalbS3 assembly.

**Figure 2.**
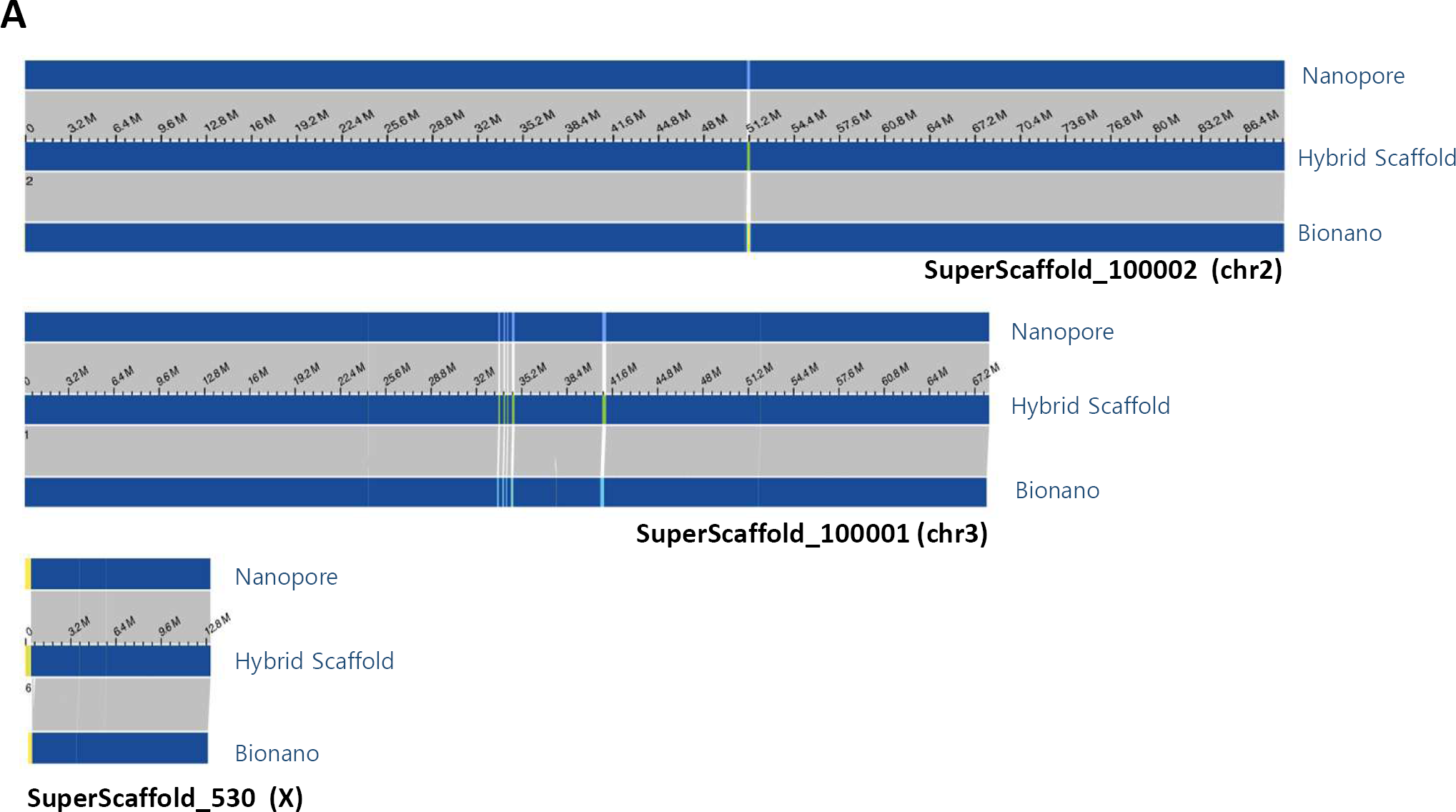
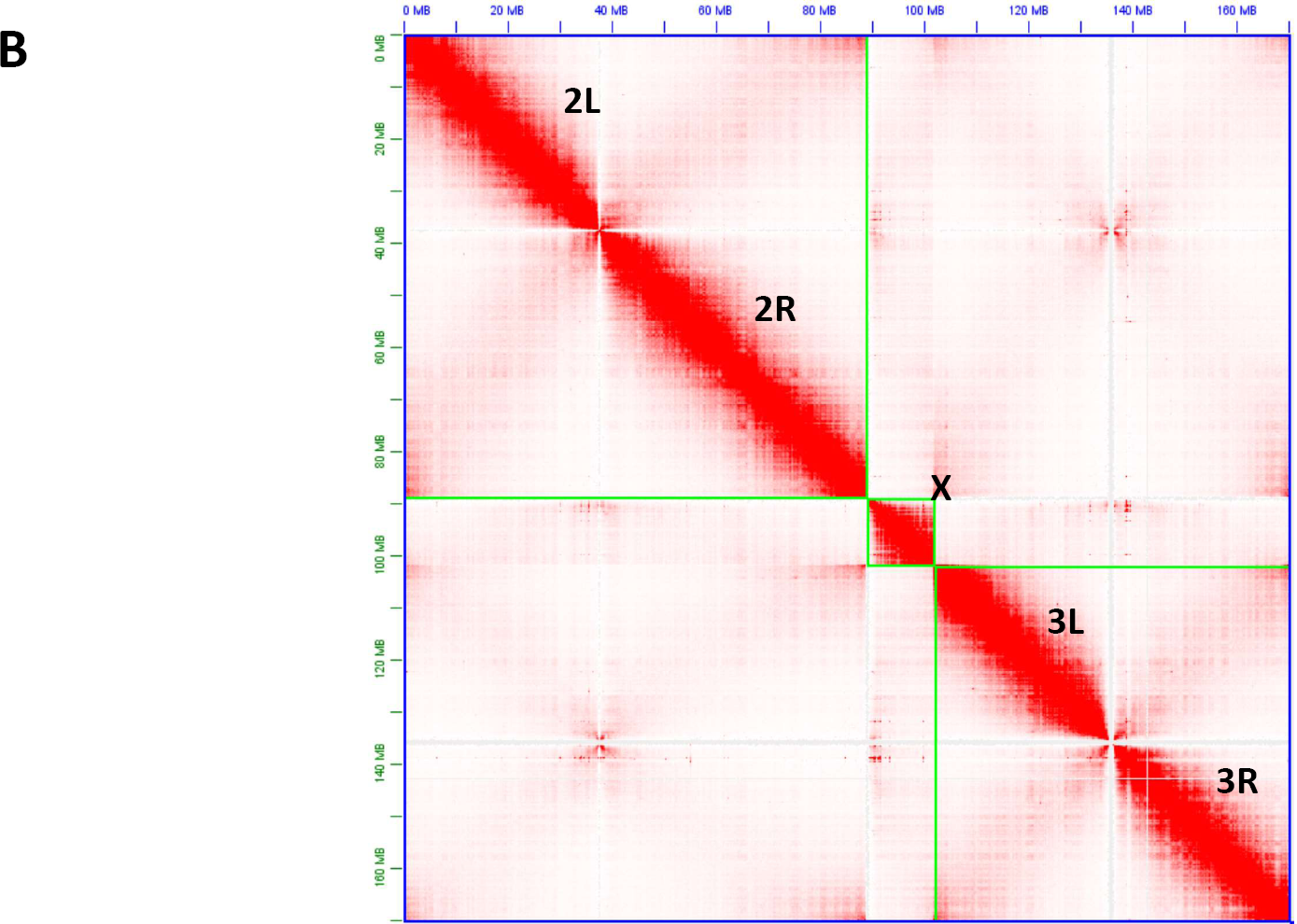
**A)** Chromosome level hybrid scaffolds produced by aligning polished Nanopore contigs to Bionano optical maps. The orientation of the hybrid scaffolds as shown in the figure from left to right: Chr2 R to L, Chr3 L to R, and X centromere to telomere. **B)** Hi-C contact matrix for the three chromosomes of the hybrid assembly. The order of the three *Anopheles albimanus* chromosomes, shown in green boxes, are 2, X, and 3. Note that the final AalbS3 assembly includes the rDNA clusters and centromeric repeats which are not shown in the figure.

### Comparison of Hi-C and Bionano scaffolding

We also produced an assembly by scaffolding the polished Canu contigs using Hi-C data, enabling informative comparisons between the two leading scaffolding technologies, Bionano and Hi-C. Using Bionano maps, the Hi-C assembly was first examined for misassembles and several chimeric mis-joins and insertions were observed that were not present in the Bionano assembly. We also assessed the quality of each assembly using physical mapping data previously used to produce the AalbS2 scaffolds [11]. Gene sequences used as probes for FISH were aligned to the Bionano and Hi-C assemblies, respectively, and their order and orientation were compared (Supplemental Table S3). We found that chromosomes X and 2 were in perfect concordance with respect to order and placement in both Hi-C and Bionano assemblies. Both Bionano and Hi-C assemblies disagreed with the FISH results in two segments on chromosome 3, indicating either chromosomal variations between samples or mistakes in FISH. Four FISH probes were not placed on the Hi-C assembly but were placed correctly in the Bionano assembly. Thus, we conclude that the overall quality of the Bionano assembly was better than the Hi-C assembly, perhaps due to the ability of the new DLS chemistry to produce ultra-long optical mapping molecules [19]. However, Bionano had difficulties assembling the tandem repeats of the centromeric region of chromosome 2 into the chromosomal scaffolds. This is likely due to an overall lack of labelling sites on the molecules. Thus, Bionano is a preferred primary scaffolding technology in this study but it should be supplemented with Hi-C for long-range scaffolding of repetitive regions that have either low or overly dense signals, similar to the strategy used in recent assemblies of plant genomes [19].

### Discovery of a novel telomeric sequence and validation by analysing the termini of long reads

We observed tandemly repeating units of 30-32bp (Figure 3A) at the ends of the chromosomes 2L, 2R, 3L, 3R and X in our AalbS3 assembly, leading us to hypothesize that these are telomeric repeat units (TRUs). As shown in Figure 3A, the 30-32 bp TRUs are very similar to each other and they form a tandemly repeated region of kilobases in length. These putative TRUs were not observed in the AalbS2 assembly as the ends of its five chromosome arms are 27-69 kb inside the chromosome arms of the AalbS3 assembly. We selected 97,921 error-corrected Oxford Nanopore reads longer than 40 kb, equivalent to ∼32x genome coverage, to validate that the AalbS3 assembly is indeed able to extend to telomeres. As shown in Figure 3B-E, reads that contain a true telomere should either start or end with the telomeric sequences while reads that contain other sequences inside the chromosomes should not all start or end with the same sequence or repeat unit. We searched the 97,921 long reads for sequences that contain any of the five TRUs (BLASTN, evalue 1e-5) and identified 149 reads. Not all 149 are expected to be telomeric sequences as the TRUs also show similarity to an 803 kb rDNA cluster scaffold and to two regions that are 77.8kb and 78.1 kb away from the 2L and X telomeres, respectively. We filtered out 65 sequences that are derived from these three regions. The remaining 84 long reads were analysed using a combination of RepeatMasker (default parameters, library being the monomers or dimers of TRUs), BLASTN (evalue 1e-5), and manual inspection. As shown in Table 2, all but one of the 84 reads either start or end with the TRU, and the only exception was likely a chimera, with ∼39kb matching 2R and ∼23 kb matching X. Thus, analysis of the long reads strongly support that the AalbS3 assembly indeed extended to the telomeres in all five chromosome arms.

**Table 2.** ONT Telomeric long reads. Only reads longer than 40 kb were analysed.

| **Chromosome Arm** | **Number of Reads beginning with TRU** | **Number of Reads terminating with TRU** | **Average Length of Reads with TRU** |
| --- | --- | --- | --- |
| **X** | 6 | 4 | 60.56 |
| **2R** | 6 | 9 | 59.86 |
| **2L** | 4 | 19 | 53.72 |
| **3R** | 17 | 7 | 54.53 |
| **3L** | 4 | 7 | 55.75 |

**Figure 3.**
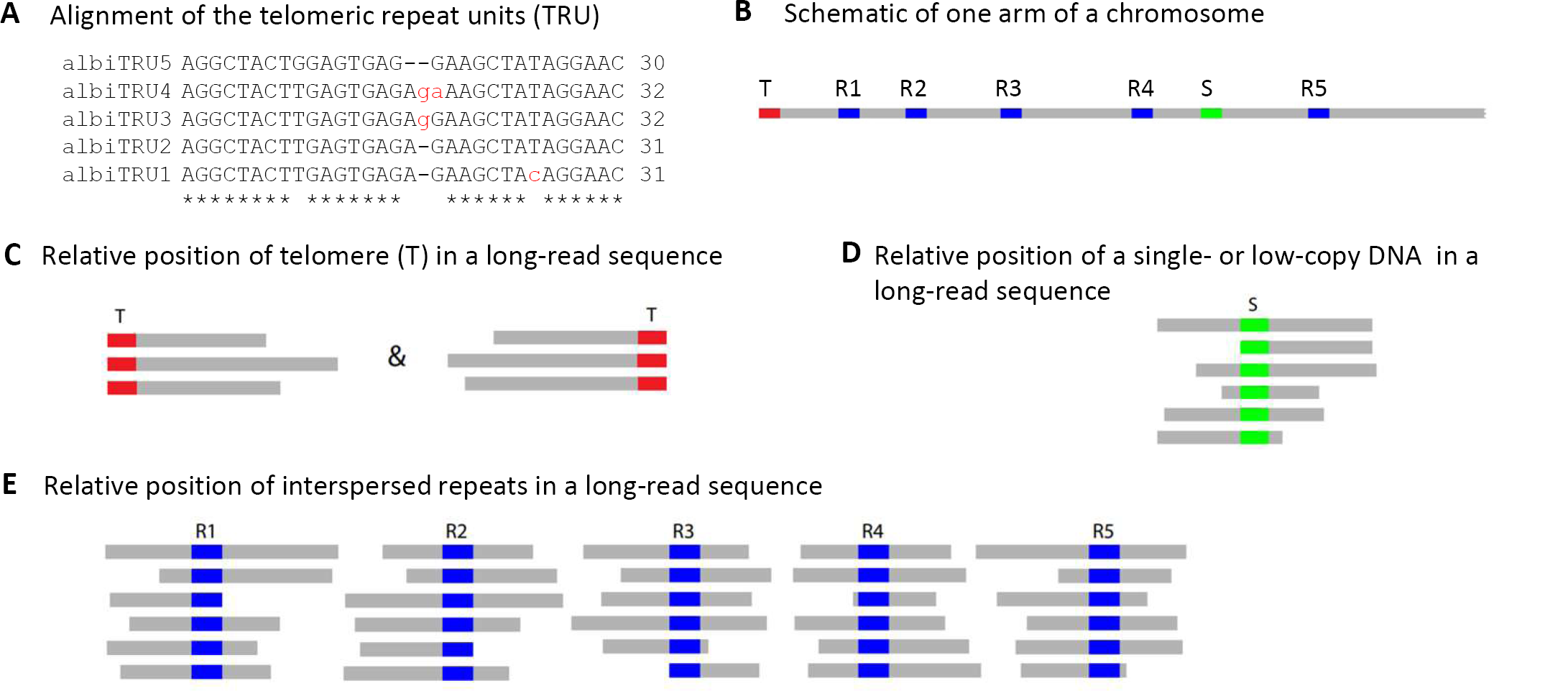
Alignments of variants of the telomeric repeat unit (TRU, panel A) and the principle underlying the strategy to verify telomere sequences using long reads (panels B-E). T: telomere, which is a tandem repeat of TRUs, or (TRU)n; S: single- or low-copy DNA sequences; R1-R5: five copies of a interspersed repeat.

### Experimental verification of the telomeric sequences

To further validate the novel 30-32 bp TRUs, we performed fluorescence *in situ* hybridization (FISH) on polytene (Figure 4) and mitotic (Figure S1) chromosomes to validate their location. A complementary oligonucleotide probe was designed based on one of the TRUs, albi_telomere1 (GTTCCTATAGCTTCTCTCACTCAAGTAGCCT). The telomeric oligonucleotide probe hybridized to all tips of chromosome arms, including 2L, 2R, 3L, 3R and X. Each arm was recognized based on the published *An. albimanus* cytogenetic map [11]. The FISH signals from telomeric repeats are present at the tips of all telomeric ends. The intensity of the signals was higher on autosomes than on the X chromosome suggesting a possibly smaller number of copies on the sex chromosome. In addition, we used a modified Bal31 exonuclease sensitivity assay [20, 21] to further validate these telomeric sequences. For this, HMW genomic DNA was digested with Bal31 to shorten ends of DNA, which was subsequently fragmented with a restriction enzyme, and detected via oligonucleotide probe hybridization by Southern Blotting (Figure 5). The telomeric probe hybridizes to Bal31-sensitive sites as indicated by progressive shortening of the detected sequences, consistent with the TRUs having a terminal position on the chromosomes.

**Figure 4.**
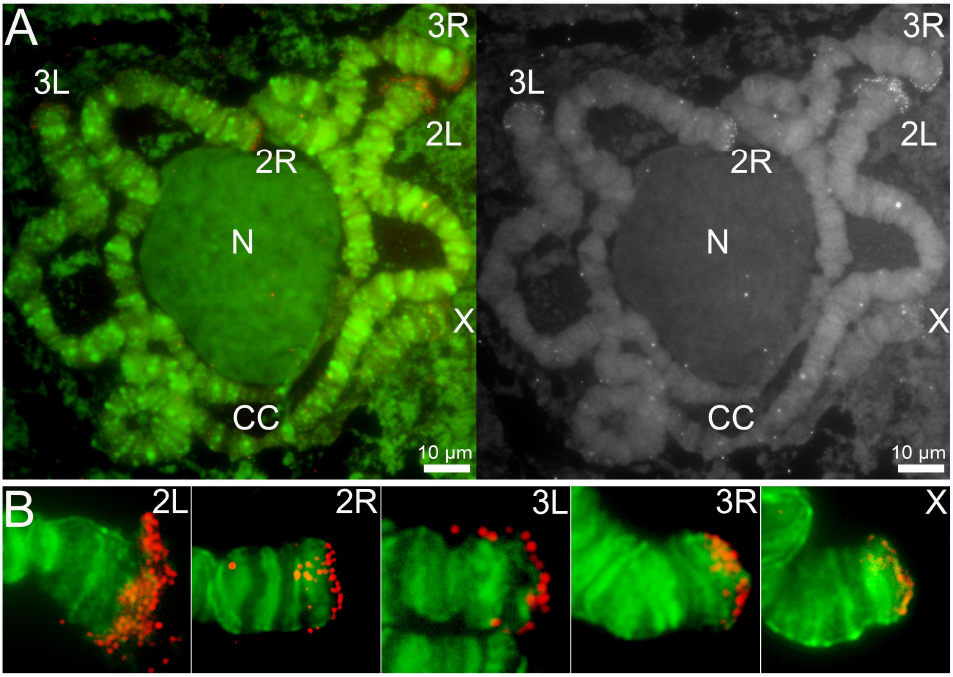
Fluorescence in situ hybridization and mapping of the *An. albimanus* telomeric oligonucleotide probe on polytene chromosomes from the 4th instar larva. (A) Left panel: Chromosomes hybridized with Cy3-labeled oligonucleotide probe (red) and counterstained with the fluorophore YOYO-1 (green). Right panel: An inverted grayscale image of the FISH results of the oligonucleotide probe. Arrows point to the *An. albimanus* telomeres. Chromosome arms are labeled as X, 2R, 2L, 3R, and 3L; the chromocenter is labeled by CC; the nucleolus is labeled by N. Scale bar – 10μm.(B). An enlarged view of representative chromosome arms hybridized with Cy3-labeled oligonucleotide probe (red).

**Figure 5.**
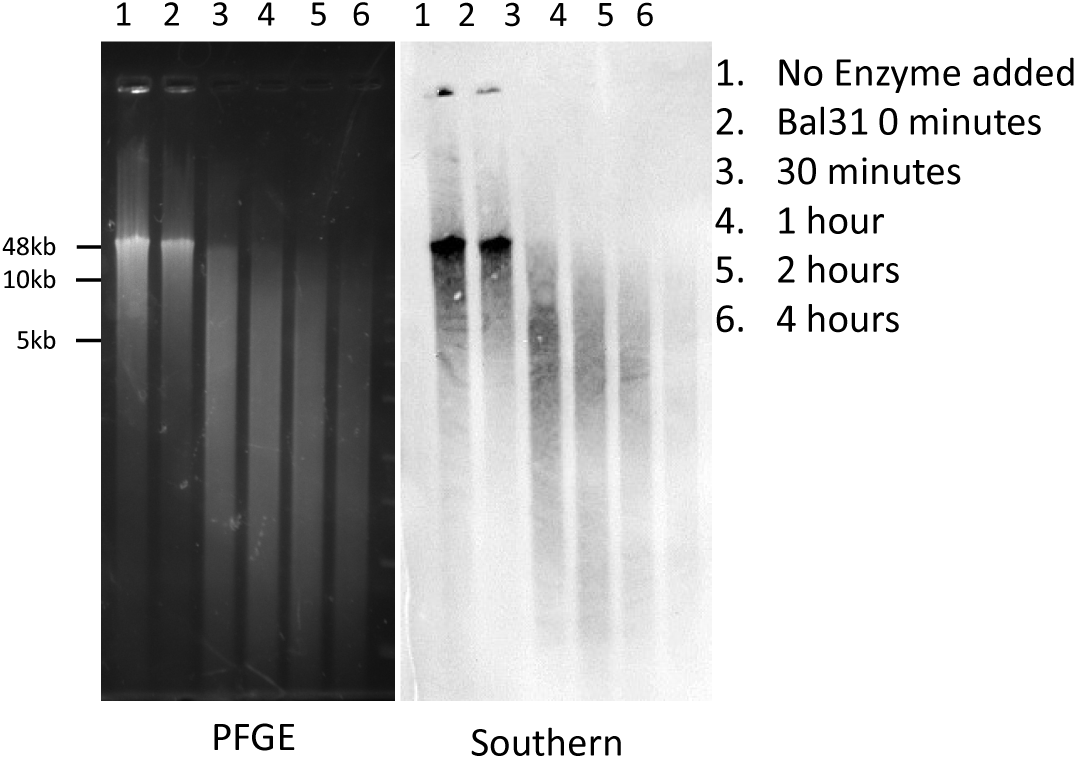
Bal31 PFGE and Southern Blot. HMW genomic DNA from *Anopheles albimanus* pupae was digested with Bal31 exonuclease at time point increments followed by restriction enzyme digestion with XbaI then separated by Pulsed-Field Gel Electrophoresis (left). The separated DNA was transferred to charged Nylon membrane via downward capillary action, hybridized with a DIG-labelled telomeric probe, and detected using chemiluminescence (right).

## DISCUSSION

In this work we have produced the AalbS3 assembly for *Anopheles albimanus*, an important vector of malaria in Central and South America. By extending to telomeres in all chromosomal arms, assembling rDNA and centromeric clusters, and recovering other repetitive sequences, the AalbS3 assembly represents a significant improvement to the previously reported AalbS2, which was one of the best *Anopheles* assemblies [1, 11]. Our experience provides insights that may inform future efforts to generate reference-quality *Anopheles* genome assemblies. First, we have shown that Oxford Nanopore sequencing is an attractive platform to generate long reads for assembling mosquito genomes. Prior to this report, only PacBio has been used as the long-read technology in place of Illumina to generate high quality mosquito assemblies [2-4]. The affordability and portability of some ONT platforms are attractive features when considering future vector sequencing projects especially for resource-limited areas or field stations. For example, we now routinely obtain >15 Gbases (60-80X coverage of an *Anopheles* genome) of long reads per ONT MinION run at a cost of approximately $600 including flow cell and library preparation. Ten Gbases of Illumina reads for polishing now cost less than $200. Second, our experience also highlights the relative effectiveness and accuracy of Hi-C and Bionano for scaffolding contigs to generate chromosome-scale assemblies. We have shown that the Bionano DLS optical mapping is a more accurate primary scaffolding method than the Hi-C method commonly used in mosquito assemblies [2, 4, 22]. However, Bionano should be supplemented with Hi-C for long-range scaffolding of repetitive regions that have either low or overly dense DLS signals [19].

An improved assembly provides a better genomic resource and can profoundly impact investigations into the molecular genetics and evolution of the species. For example, information on previously uncharacterized centromeric repeats will facilitate chromosomal analysis during mitosis and meiosis. Furthermore, recovering a large number of repeats will inform analysis of genome structure and repeat-associated small RNAs. Most importantly, we discovered novel telomeric repeats present at chromosomal termini. These differ in both sequence and structure from telomeric repeats reported in other dipteran species, which may indicate a novel telomere synthesis or maintenance mechanism and provide new evolutionary insights. The approach we used to validate the telomeres took advantage of the ONT long reads as TRU-containing long reads should either begin or end with TRU sequences. This method could be generally applicable for the discovery and validation of telomeres in diverse organisms.

There are generally three known mechanisms that maintain telomere length in eukaryotes. The most common mechanism used by many organisms including humans, employs a telomerase consisting of a reverse transcriptase and an RNA template to extend short tandem repeats at chromosome ends [23]. The second, described in *Drosophila*, involves assembly of HeT-A and TART retrotransposon arrays at chromosomal termini [24-26]. The third mechanism, as described in *An. gambiae* through a serendipitous discovery of a transgene inserted into the telomeric regions, possibly relies on unequal crossover of telomeric repeats that are hundreds of bases long [27, 28]. Contrasting the 820bp satellite repeat units reported in *An. gambiae*, here we describe much smaller 30-32bp TRUs at the chromosome ends of *An. albimanus*. BLAST analysis indicates that the *An. albimanus* TRUs do not resemble the *An. gambiae* telomeric satellite nor the *Drosophila* telomeric retrotransposons. It is not yet clear how telomeres are maintained by these novel TRUs in *An. albimanus*. What is also unknown is whether telomeric lengths are correlated with the female life span in mosquitoes, which is a life history trait relevant to disease transmission as only older females are responsible for pathogen transmission [29, 30].

## MATERIALS AND METHODS

### Mosquito strain

The STELCA strain of *An. albimanus* was originally colonized from a population in El Salvador and deposited at the Malaria Research and Reference Reagent Resource (MR4) at Biodefense and Emerging Infections Research Resources Repository (BEI) under catalog number MRA-126. A colony of this strain was established and maintained in the insectary of the Fralin Life Science Institute at Virginia Tech. All stages were reared in a growth chamber at 27°C with a 12-hour light cycle.

### Mosquito mating scheme and sample collection

A single male mosquito and five virgin female mosquitoes were allowed to mate for 4 days in a 16 oz soup cup then given a blood meal using defibrinated sheep blood using an artificial blood-feeder. After 72 hours, eggs were collected from each female separately and reared under normal conditions. All F_1_ pupae from each family were sorted by sex, collected, and immediately frozen in liquid nitrogen and stored in −80° C. The largest family was chosen for sequencing purposes and the F_0_ father and the F_0_ mother were collected separately, flash frozen in liquid nitrogen, and stored at −80° C. This sample collection scheme is depicted in Figure 1.

### DNA isolation

Genomic DNA was isolated from 20 female F_1_ pupae following a modified Qiagen Genomic Tip DNA Isolation kit (Qiagen Cat No. 10243 and 19060) protocol. For this, pupae were immediately transferred from −80° C into a 50mL conical tube containing pre-prepared lysis solution consisting of 9.5mL Buffer G2 and 19μL RNase A (Qiagen Cat No. 19101). The pupae were then homogenized using a Dremel motorized homogenizer for approximately 30 seconds on the lowest speed. Next, >300mAU (500μL of >600 mAU/ml, solution) Proteinase K (Qiagen Cat No. 19131) was added to the sample and incubated at 55° C for 3 hours. The homogenate was then transferred into a 15mL conical tube and centrifuged at 5000x G for 15 minutes at 4° C to remove debris. Following this, DNA was extracted following the standard Qiagen Genomic Tip protocols. The purity, approximate size, and concentration of the DNA were tested using a nanodrop spectrophotometer, 0.5% agarose gel electrophoresis, and Qubit dsDNA assay, respectively.

### Oxford Nanopore sequencing and base calling

Approximately 1μg of DNA was used to generate a sequencing library according to the protocol provided for the SQK-LSK109 library preparation kit from Oxford Nanopore. After the DNA repair and end prep and adapter ligation steps, SPRIselect bead suspension (Beckman Coulter Cat No. B23318) was used to remove short fragments and free adapters. Qubit dsDNA assay was used to quantify DNA and approximately 300-400 ng of DNA library was loaded onto a MinION flow cell. Base calling was performed using albacore with the default filtering setting of Qscore >7 [31] (BioProject Number PRJNA622927, BioSample SAMN14582895).

### Illumina sequencing library preparation and sequencing

Genomic DNA was isolated from F_0_ mother and F_0_ father using the QiaAMP DNA micro kit (Qiagen Cat No. 56304). Approximately 300 ng of genomic DNA was used to prepare DNA sequencing libraries for each parent following the protocol of NEBNext Ultra II FS DNA Library Prep Kit for Illumina (NEB Cat No. #E7805S/L). The libraries were sent to Novogene (https://en.novogene.com/) for sequencing. More than 50 Gb of 2×150 bp reads were obtained from the father and mother, respectively (BioProject Number PRJNA622927, BioSamples SAMN14582897 and SAMN14582898, respectively).

### Contig assembly and polishing

Quality-filtered sequences longer than 2 kbps were assembled with Canu [14] using the cascades high-performance computer at the Advanced Research Computing facility of Virginia Tech. This generated a 197.39 Mbps genome consisting of 660 contigs with an N50 of 13.7 Mbps. These contigs were polished with Pilon [15] for four rounds using parental Illumina short reads. The completeness of the assembly was measured using a Benchmarking Universal Single Copy Orthologs (BUSCO) test [17, 18]

### Hi-C scaffolding

Hi-C Illumina reads were available from a multi-species Hi-C analysis project (PRJNA615337) that included 2 replicates of *Anopheles albimanus* mixed embryo samples (SAMN14451359). Hi-C reads were aligned to either the polished Canu contigs for independent scaffolding (Table S3) or to the Bionano scaffolds for validation (Figure 2B). 3D-DNA pipeline was employed to assemble the *An. albimanus* genome *de novo* using the generated Hi-C data set. Misassemblies were identified and fixed manually using assembling mode in Juicebox software[32]. The *An. albimanus* physical genome map was used to assess the assemblies (Table S3) [11].

### Ultra-high molecular weight (uHMW) nuclear DNA isolation and Bionano Mapping

Ultra-high molecular weight (uHMW) nuclear DNA was isolated from approximately 40 pooled male siblings (Figure 1) using a modification of the Bionano Prep™ Animal Soft Tissue DNA Isolation protocol (bionanogenomics.com; document 30077). Flash-frozen pupae were homogenized with a chilled Dounce grinder in the presence of ice-cold Bionano Prep™ homogenization buffer. The sample was then filtered through a single 100um cell strainer, mixed with an equal part of cold 200-proof ethanol, and incubated for one hour at room temperature. Nuclei and debris were pelleted by centrifugation at 1,500xg for 5 minutes at 4C, followed by four wash-centrifugation cycles with homogenization buffer. The resulting pellet was embedded in three 90-ul low-melting-point agarose plugs and treated with both proteinase K and RNaseA as per manufacturer’s recommendations. Free ultra-high molecular weight nuclear DNA was recovered by melting the plugs, digesting with agarase and dialyzing against TE buffer. Data collection for optical mapping was performed in a Bionano Saphyr platform running a sample prepared according to the Direct Label and Stain (DLS) process (Bionano Genomics Cat.80005) following manufacturer’s protocols with some modifications. Approximately 500ng uHMW nDNA was incubated for 2:20 h at 37 °C, followed by 20 min at 70 °C in the presence of DLE-1 Enzyme, DL-Green and DLE-1 Buffer. The labelling reaction was killed by adding proteinase K and incubate at 50C for 1hr, followed by clean-up of the unincorporated DL-Green label. Labeled, cleaned-up DNA was then combined with a Flow/DTT buffer as per Bionano Genomics specifications and incubated overnight at 4 °C. After quantification, DNA was stained by adding Bionano DNA Stain to a final concentration of 1 microliter per 0.1 microgram of final DNA, loaded into a Saphyr chip flowcell. Molecules were stretched, separated, imaged and digitized using software installed in a Bionano Genomics Saphyr System and server according to the manufacturer’s recommendations (https://bionanogenomics.com/support-page/saphyr-system/). Molecules were automatically filtered with a minimum length of 150 kb and a minimum of 9 labels, resulting in a 342-Gbp subset of 1,014,877 molecules with a N50 of 373 kbp and average label density of 9.85 per 100 kbp. The molecules were assembled into maps by the Bionano Solve Version 3.3, RefAligner Version 7989 and Pipeline Version 7981 software. The used parameters were “non-haplotype without extend and split”, no CMPR cuts, and 200 Mbp expected genome size. The resulting assembly included 29 maps with an N50 of 51 Mbp and a total combined length of 223 Mbp. Bionano (BNG) maps were aligned to the described sequence assembly to generate hybrid scaffolds using the Bionano Solve V3.3 suite. The alignment produced only 8 scaffolds, with a total scaffold length of 184.23 Mbp, including a 11.8 Mbp map with little corresponding sequence from the female sequence assembly. Sequence was manually curated to trim sequence overlaps, remove secondary contigs due to heterozygosity, break up mis-assemblies and ultimately validate final sequence.

### Repeat analysis

The AalbS3 assembly was used to uncover repeat sequences using RepeatModeler (http://www.repeatmasker.org/RepeatModeler/) with default settings. The repeat library generated by RepeatModeler was then used to mask either the AalbS2 or AalbS3 assembly, by using RepeatMasker (http://www.repeatmasker.org) with default settings, for comparison of repeat content.

### Chromosome preparation and fluorescence *in situ* hybridization

Polytene chromosome and mitotic chromosome preparations were made as previously described [33]. Salivary glands were dissected from one 4th instar larvae, which was stored in Carnoy’s solution (Methanol: Glacial acetic acid = 3:1), and used for one polytene chromosome preparation. Isolated salivary glands were bathed in a drop of 50% propionic acid for 5 min and squashed under a 22 × 22 mm cover slip. Mitotic chromosome preparations were made from leg and wing imaginal discs of early 4th instar larvae. A fresh drop of hypotonic solution was added to the preparation of imaginal discs for 10 min, followed by fixation in a drop of modified Carnoy’s solution (ethanol:glacial acetic acid, 3:1) for 1 min. Next, a drop of freshly prepared 50% propionic acid was added, and the imaginal discs were covered with a 22 × 22 mm coverslip. The quality of chromosomal preparation was assessed with an Olympus CX41 phase-contrast microscope (Olympus America Inc., Melville, NY). High-quality chromosome preparations were flash frozen in liquid nitrogen, coverslips were quickly removed by a razor blade and preparations were immediately placed in cold 50% ethanol. After that, preparations were dehydrated in an ethanol series (50%, 70%, 90%, and 100%), air-dried and stored at room temperature (RT) until the use for FISH. To prepare the probe, an oligonucleotide was designed based on sequences of the telomeric satellite albi_telomere1: GTTCCTATAGCTTCTCTCACTCAAGTAGCCT and labelled with 3’-end-Cyanine3 fluorochrome (Sigma-Aldrich, St. Louis, MO, USA) The sequence of the oligonucleotide probe is AGGCTACTTGAGTGAGAGAAGCTATAGGAAC [Cyanine3]. FISH was performed as previously described [34, 35] with modifications. Briefly, slides with good chromosome preparations were washed in 1×phosphate-buffered saline (PBS) for 20 min and fixed in 3.7% formaldehyde for 1 min at RT. Slides were then washed in 1×PBS briefly and dehydrated in a series of 70%, 90%, and 100% ethanol for 5 min at RT. Then, 10 µl of 100 μM oligonucleotide probes diluted in the hybridization buffer, including 1200 μl deionized formamide, 0.2 g Dextran Sulphate, 120 μl 20×SSC and 500 μl H 2 O, were added to the preparations. After heating at 73 °C for 5 min, slides were incubated at 37 °C overnight. After washing in 1×SSC at 60°C for 5 min and 4×SSC/NP40 solution at 37°C for 10 min, slides were briefly washed in 1×PBS and incubated in YOYO-1 for 10 min at RT. After rinsing in 1×PBS, preparations were counterstained with an antifade solution (Life Technologies, Carlsbad, CA, USA) and kept in the dark for at least 2 hours before visualization using a ZEISS Axio Imager 2 fluorescent microscope (Zeiss, Oberkochen, Germany) with a connected Axiocam 506 mono digital camera (Zeiss, Oberkochen, Germany).

### Bal31 Sensitivity Assay

HMW genomic DNA was extracted from 50 *Anopheles albimanus* following an SDS-based method as previously described [36]. The Bal31 Sensitivity Assay was performed based on the protocol of [20, 21] with several modifications. Briefly, 2 µg of HMW genomic DNA was treated with Bal31 exonuclease for the prescribed amount of time (0, 30, 60, 120, and 240 minutes) followed by inactivation by the addition of ethylene glycol tetraacetic acid (EGTA) to a final concentration of 20 mM and incubated at 65 C for 5 minutes. The digested DNA was recovered using a phenol-chloroform extraction and ethanol precipitation as described above. The recovered DNA was treated with XbaI for two hours at 37 C and inactivated by heating at 65 C for 20 minutes. Following this, 20 µL of each sample was analysed by Pulsed-Field Gel Electrophoresis (PFGE) for 8 hours at 14 C (6 V/cm, 10-50 seconds switch time). PFGE-separated DNA was transferred onto Hybond-N+ charged nylon membrane by downward capillary transfer. Hybridization of the digoxegenin (DIG)-labelled 31bp oligonucleotide probe and detection was carried out following the DIG High-Prime Labelling and Detection Starter Kit (Roche SKU 11745832910).

100 µl of extraction buffer (200 mM Tris-HCl, pH 8.0; 0.5% Sodium dodecyl sulphate; 250 mM NaCl; 50mM EDTA)

## Data Availability

1. The AalbS3 *Anopheles albimanus* genome assembly: NCBI Genome database https://www.ncbi.nlm.nih.gov/genome BioProject PRJNA622927.
2. Raw data deposited in NCBI Sequencing Read Archive (SRA) https://www.ncbi.nlm.nih.gov/sra/ under PRJNA622927: F_1_ females ONT SAMN14582895, F_0_ father SAMN14582897, F_0_ mother SAMN14582898
3. Bionano molecules are deposited in NCBI Sequencing Read Archive (SRA) https://www.ncbi.nlm.nih.gov/sra/under PRJNA622927: F_1_ males SAMN14582896.
4. HiC sequencing reads are deposited in NCBI Sequencing Read Archive (SRA) https://www.ncbi.nlm.nih.gov/sra/ under PRJNA615337: 2 replicates *Anopheles albimanus* SAMN14451359.

The AalbS3 assembly FASTA, the repeat library, and the 84 telomeric ONT reads were uploaded and are available through the BioRxiv submission portal as Supplemental Files. The authors affirm that all data necessary for confirming the conclusions of the article are present within the article text, figures, and tables.

## Acknowledgement

This work is supported by NIH grants AI133571 to ZT, AI135298 to IVS, and the Virginia Agriculture Experimental Station. AC is supported by a fellowship from the Robert Wood Johnson Foundation. We thank the Advanced Research Community at Virginia Tech for access to high performance computer clusters used for genome assembly.

## Supplemental Tables

**Table S1.** Repeat comparison of AalbS2 and AalbS3.

|  | **AalbS2** | **AalbS3** |
| --- | --- | --- |
| **Total Repeat Content** | 2.20% | 5.06% |
| SINEs | 0.01% | 0.01% |
| LINEs | 0.64% | 0.83% |
| LTR retrotransposon | 0.16% | 0.53% |
| DNA elements | 0.55% | 1.31% |
| Unclassified | 0.84% | 2.38% |
| Simple repeats | 0.26% | 0.18% |

**Table S2.** BUSCO test comparison of AalbS2 and AalbS3.

| BUSCO<br>Classification | AalbS2 |  | AalbS3 |  |
| --- | --- | --- | --- | --- |
|  | Group<br>hits | Percentage | Group<br>hits | Percentage |
| Complete | 1055 | 98.9 | 1055 | 99.0 |
| Complete and<br>single-copy | 1046 | 98.1 | 1047 | 98.2 |
| Complete and<br>duplicated | 9 | 0.8 | 8 | 0.8 |
| Fragmented | 3 | 0.3 | 4 | 0.4 |
| Missing | 8 | 0.8 | 7 | 0.6 |

**Table S3.**
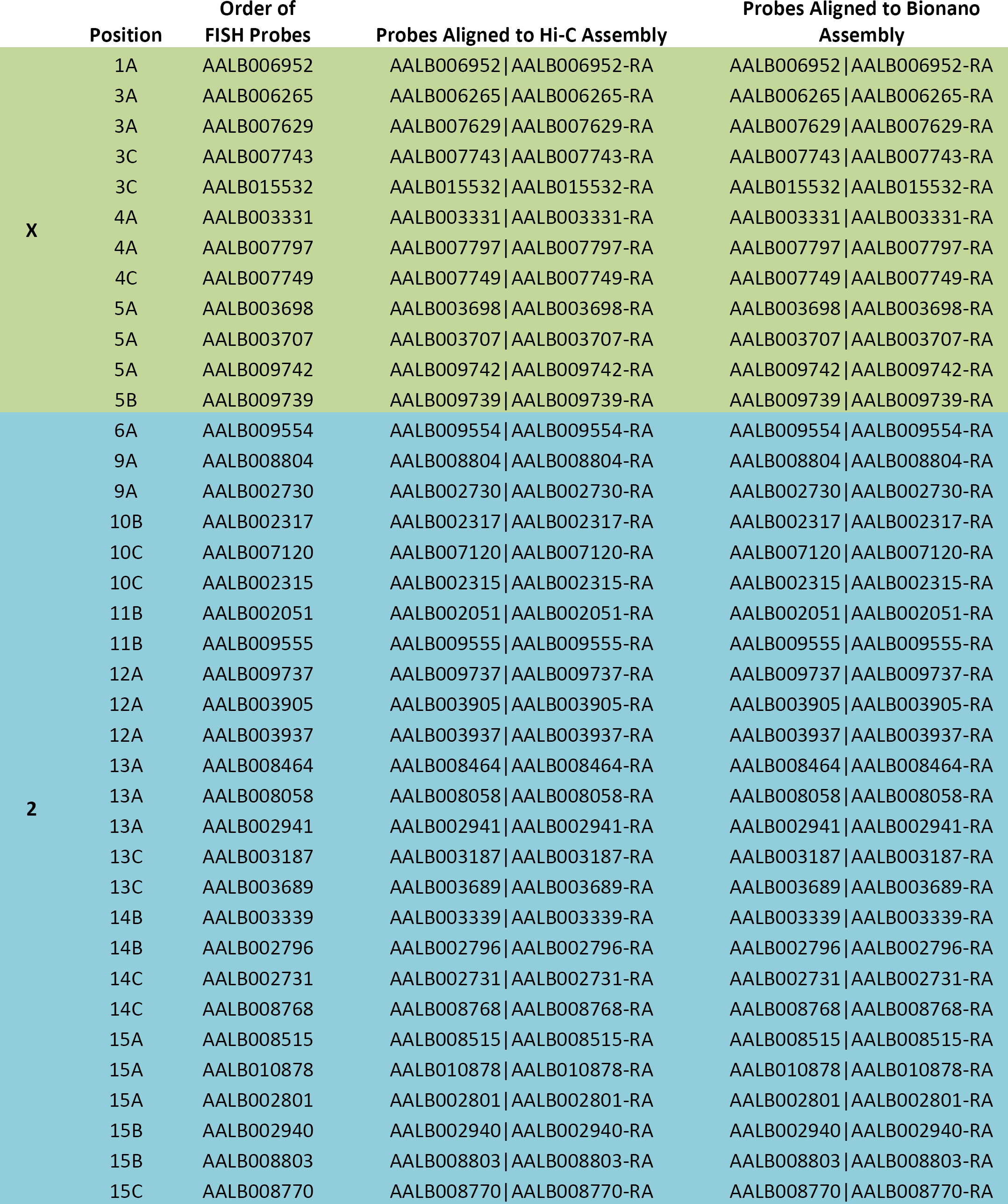
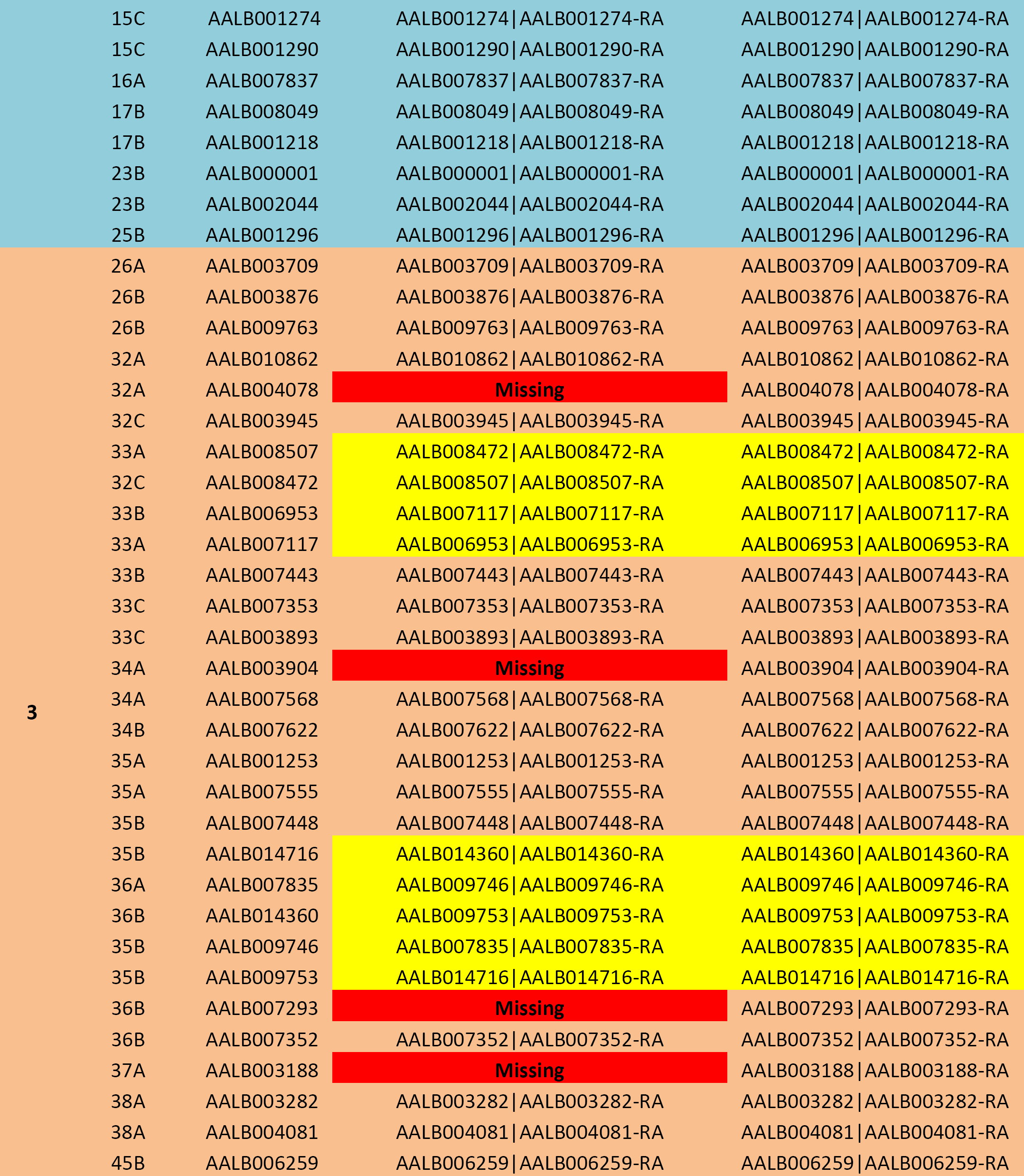
FISH BLAST Analysis.

## List of supplemental materials

Figure S1

Tables S1-S3

Supplemental Data File 1: AalbS3 assembly FASTA (BioProject PRJNA622927)

Supplemental Data File 2: Repeat library

Supplemental Data File 3: The 84 ONT long reads that begin or end with TRUs plus one chimeric read

---

**Figure S1.**
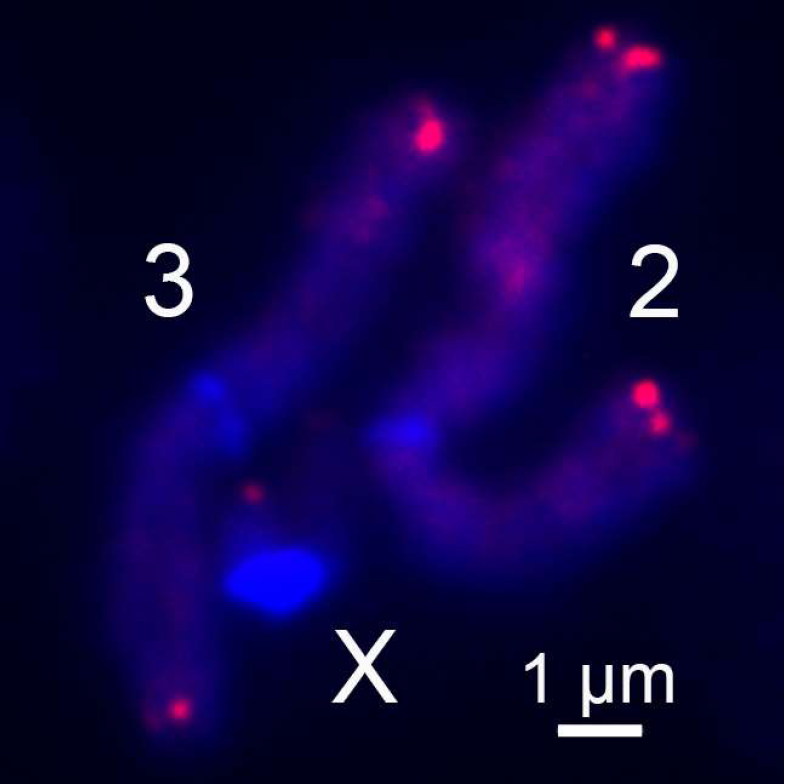
Fluorescence *in situ* hybridization and mapping of the *An. albimanus* telomeric oligonucleotide probe on mitotic chromosomes from the 4th instar larva. Chromosomes hybridized with Cy3-labeled oligonucleotide probe (red) and counterstained with the fluorophore DAPI (blue). Chromosome arms are labeled as X, 2, and 3; Scale bar – 1 μm.

## Notes

### Competing Interest Statement

The authors have declared no competing interest.

### Summary of Updates

Updated supplemental files and Data Availability section; minor changes to Table 1.

